# BBmix: a Bayesian Beta-Binomial mixture model for accurate genotyping from RNA-sequencing

**DOI:** 10.1101/2022.12.02.518817

**Authors:** Elena Vigorito, Anne Barton, Costantino Pitzalis, Myles J. Lewis, Chris Wallace

## Abstract

**Motivation:** While many pipelines have been developed for calling genotypes using RNA-sequencing data, they all have adapted DNA genotype callers that do not model biases specific to RNA-sequencing such as reference panel bias or allele specific expression.

**Results:** Here, we present BBmix, a Bayesian Beta-Binomial mixture model that first learns the expected distribution of read counts for each genotype, and then deploys those learned parameters to call genotypes probabilistically. We benchmarked our model on a wide variety of datasets and showed that our method generally performed better than competitors, mainly due to an increase of up to 1.4% in the accuracy of heterozygous calls. Moreover, BBmix can be easily incorporated into standard pipelines for calling genotypes. We further show that parameters are generally transferable within datasets, such that a single learning run of less than one hour is sufficient to call genotypes in a large number of samples.

**Availability:** We implemented BBmix as an R package that is available for free under a GPL-2 licence at https://gitlab.com/evigorito/bbmix and accompanying pipeline at https://gitlab.com/evigorito/bbmix_pipeline.

## Introduction

RNA-sequencing (RNA-seq) is an established and popular technique with many applications including transcript quantification and detection of alternative splicing among others. More recently, a growing number of studies have developed and tested pipelines to use RNA-seq for variant and genotype calling (Rogier *et al*., 2018), (Quinn *et al*., 2013), (Adetunji *et al*., 2019), (Brouard *et al*., 2019) including somatic variant detection and identification of cancer drivers (Akutagawa *et al*., 2022). Applications include calling variants when DNA genotypes are unavailable (Wang *et al*., 2021), studying *cis*-regulated genes by analyzing allele specific expression (ASE) or conducting genome-wide association studies (GWAS) in non-model species, since whole genome sequencing is expensive and exome sequencing tools may be unavailable (Jehl *et al*., 2021), (Rogier *et al*., 2018). Genotype calling using RNA-seq is often more challenging than DNA-seq: coverage can be highly variable between genes, many reads map across splice junctions making correct alignment more difficult, and cis-acting eQTLs may create unequal expression between chromosomes. Despite this, for many applications, accuracy is paramount: for example in ASE, miscalling a homozygous genotype as heterozygous will create a false signal of allelic imbalance.

RNA-seq genotyping pipelines employ RNA-seq based aligners and quality control steps for filtering out potentially erroneous calls, but use the same statistical models as for DNA-sequencing. While adopting these methods provides a convenient way to call variants from RNA-seq, there are sources of variation in RNA-seq data that are not accounted for. In DNA-seq, in an ideal world, out of n reads covering a SNP, we expect 0, n/2 and n to contain the alternative allele for genotypes 0, 1 and 2. Sampling variation and reference mapping bias may mean the observed heterozygote genotype count deviates from n/2 particularly when n is small, and the homozygote observations may deviate minorly from 0 and n due to sequencing or mapping errors. However, when n is large, the genotypes are expected to be clearly distinguishable. In RNA-seq, the heterozygote observation may show much greater variance, because heterozygosity at local regulatory sequences may cause over-amplification of the allele on one chromosome over the allele on the other, and this effect will vary between samples as genotypes at the regulatory sequences vary. This may push the heterozygote observation nearer to 0 or n, making a miscall as a homozygote more likely, particularly when n is small.

In this work, we wanted to test whether directly modelling this additional variation in RNA-seq data could improve the quality of the genotype calls. We developed BBmix, a two-step method based on first modelling the genotype-specific read counts using beta binomial distributions and then using these to infer genotype posterior probabilities. ir We benchmarked our method using high confidence calls from the Genome in a Bottle consortium (https://www.nist.gov/programs-projects/genome-bottle), the subset of GBR samples from the GEUVADIS (Genetic European Variation in Disease) project (McVean *et al*., 2012), for which RNA-seq data and high quality genotype calls are publicly available and samples from the Pathobiology of Early Arthritis Cohort (PEAC) cohort (Lewis *et al*., 2019) for a “real data” case study. This is a cohort of 82 adult treatment-naive rheumatoid arthritis patients for whom RNA-seq from synovial tissue samples and DNA-microarray based genotypes are available. We found that our method generally performed better than competitors and can be easily incorporated into standard pipelines for calling variants.

## Methods

### BBmix statistical model

We model the number of reads at the alternative allele at each SNP as a mixture of three beta-binomial distributions underlying the three possible genotypes, conditional on the total number of reads overlapping the SNP. To allow for sequencing or alignment errors as well as reference mapping bias, we allowed both the mean and variance parameters of each distribution to be learned from the data. The parameters of these distributions are shared between SNPs in a Bayesian model allowing the parameters to be learnt from many SNPs jointly. To maximise computational efficiency, we train the model on a random subset of SNPs and use the learnt parameters to call genotype probabilities across all SNPs. Details of the model are presented in **Supplementary Note 1**.

### Datasets

We use RNA-seq from lymphoblastic cells lines derived from NA12878 and 13 additional samples (study E-MTAB-1883) with the high confidence genotype calls from the Genome in a Bottle consortium for sample NA12878 used as a gold standard. Note that only non-reference homozygote genotypes are available for this data source. We also selected a subsample of 86 individuals of European ancestry coded as GBR from the GEUVADIS for which genotypes and RNA-seq data are publicly available and genotypes and RNA-seq from the PEAC study. Details for data sources both for genotypes and RNA-seq can be found in **Supplementary Note 2**.

### RNA-seq preprocessing

We applied the GATK pipeline for RNA-seq short variant discovery (SNPs + Indels, https://gatk.broadinstitute.org/hc/en-us/articles/360035531192-RNAseq-short-variant-discovery-SNPs-Indels-) for read alignment and data cleanup (removal of duplicates and base quality recalibration) before calling variants. Details on the implementation are described in **Supplementary Note 3**.

### Comparison to alternative methods

We validated BBmix against three other popular methods: HaplotypeCaller (Brouard *et al*., 2019), Mpileup from Bcftools (Li, 2011) and FreeBayes (Garrison and Marth, 2012) which have been applied for RNA-seq genotyping (Wang *et al*., 2021; Rogier *et al*., 2018; Adetunji *et al*., 2019; Quinn *et al*., 2013). We also compared BBmix with a naive method based on applying thresholds on the proportion of alternative alleles; we referred to this method as “Count threshold”. The alignment output from the GATK pipeline was used as input for calling genotypes with BBmix, Count threshold, Haplotypecaller and Mpileup/bcftools. For FreeBayes the base quality recalibration step was excluded as it is not recommended (https://github.com/freebayes/freebayes). Each method was run with default parameters. Details of the implementation are in **Supplementary Note 4**. We compared hard calls for exonic variants in the 1000 Genome project with allele frequency higher than 0.01 and excluded regions of the genome with known mapping bias against the corresponding gold standard (**Supplementary Note 4)**. For the PEAC gold standard for which probabilistic genotype calls were available, we selected the subset for which genotypes were called with probability 1.

### Code availability

The R package with the code for genotyping using bbmix can be found at https://gitlab.com/evigorito/bbmix. The pipeline for preparing the inputs for running bbmix can be found at https://gitlab.com/evigorito/bbmix_pipeline. The pipeline and code used for preparing the manuscript is at https://gitlab.com/evigorito/bbmixpaper_pipeline.

## Results

### Determining the number of SNPs for training the Beta-Binomial mixture model

We first evaluated the effect of the number of SNPs used to train the Beta-Binomial mixture (BBmix) model on the accuracy of genotype calls. To this end, **three sets** of 500, 1,000, 2,000, 3,000, 4,000, 5,000 or 10,000 SNPs with at least 10 mapping reads (**21 sets** in total) were randomly selected from the Genome in a bottle sample NA12878. We fit the BBmix model independently on each subset (Methods). As expected, increasing the number of SNPs resulted in a narrower posterior distribution of the mean parameter of each component (**Fig. 1a)**, while the posterior mean for heterozygous calls departed from 0.5, suggestive of over-representation of the reference allele consistent with reference mapping bias (**Fig. 1a medium panel)**. Increasing the number of training SNPs also resulted in overall small gains in concordance between genotype dosages estimated by BBmix and the high confidence calls available from the Genome in a bottle consortium until a plateau was reached at about 4,000 SNPs (**Fig. 1b**). For subsequent analysis we trained our model with 1000 SNPs, which we considered a good trade-off between concordance and computational time (mean time=46 minutes, **Fig. 1c)**.

**Figure 1.**
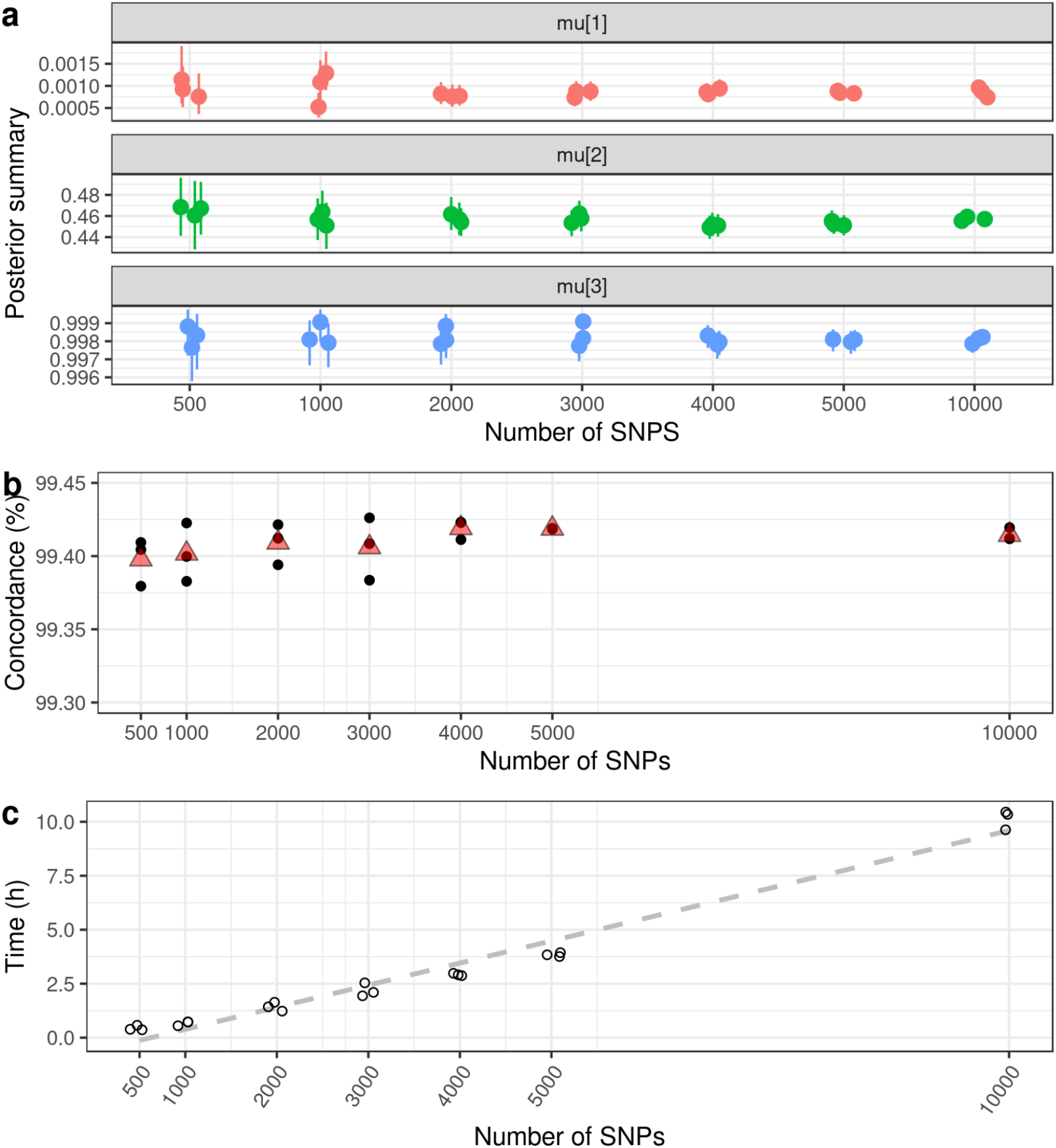
Training the Beta-Binomial mixture (BBmix) model with different numbers of SNPs. A random sample of RNA-seq reads for the indicated number of SNPs with at least 10 supportive reads were used to train the BBmix model. The procedure was repeated 3 times. (a) Mean and 95% credible interval for the posterior distribution for each of the 3 components of the mean parameter (mu[1-3]). (b) Concordance of calls (y-axis) based on the expected genotypes (dosages) called with the parameters learnt from the BBmix model trained with the indicated number of SNPs (x-axis). The black points correspond to each replicate and the red triangle corresponds to the mean. (c) Running time (h) for fitting the BBmix model for the indicated number of SNPs. The average time when training with 500 SNPs was 25 minutes; for 1,000 SNPs was 46 minutes; for 2,000 83 minutes; for 3,000 128 minutes; for 4,000 2.9 hours; for 5,000 SNPs 4 hours and for 10,000 SNPs 10 hours. The dashed line corresponds to the linear regression line.

Some analyses require hard calls instead of genotype dosages. To define hard calls we assigned the most likely genotype when its probability is above a predefined threshold. We tested thresholds of 0.90, 0.95 and 0.99 for sample N12878 and chose to favour accuracy over number of calls. We selected 0.99 as at depth 10 the concordance was 99.82% (0.1% increase relative to 0.9) while the loss in the number of calls was 0.15% (9303 total calls, **Supp. Fig. 2**). In subsequent analyses, when applying the BBmix model on hard calls we selected 0.99 as the probability threshold.

### Benchmarking the BBmix model

We validated the BBmix model against three other popular methods: HaplotypeCaller (Brouard *et al*., 2019), Mpileup from Bcftools (Li, 2011) and FreeBayes (Garrison and Marth, 2012) which have been applied for RNA-seq genotyping (Wang *et al*., 2021; Rogier *et al*., 2018; Adetunji *et al*., 2019; Quinn *et al*., 2013). We also compared BBmix with a naive method based on applying thresholds on the proportion of alternative alleles; we referred to this method as “Count threshold”. We applied these methods to three datasets: the highly curated genome in a bottle sample, a subset of the GEUVADIS samples with high quality genotypes and samples from the PEAC study which were genotyped by DNA-microarrays.

We first assessed the accuracy of the calls stratified by genotype across the different datasets. For this analysis we hard called genotypes for SNPs with at least 10 supporting reads, which we considered as a good trade-off between coverage and accuracy (**Supp. Fig. 2-4)**. Homozygous reference calls tend to be the most accurate (> 99.6%) with the lowest variance across methods (**Fig. 2**), while heterozygous calls are generally more prone to error (accuracy < 99.0% in GEUVADIS and PEAC) and showed the highest variability across methods (**Fig. 2**). Overall, BBmix was generally superior to competitor methods, with the main gains in the accuracy of heterozygous calls (**Fig. 2**).

**Figure 2.**
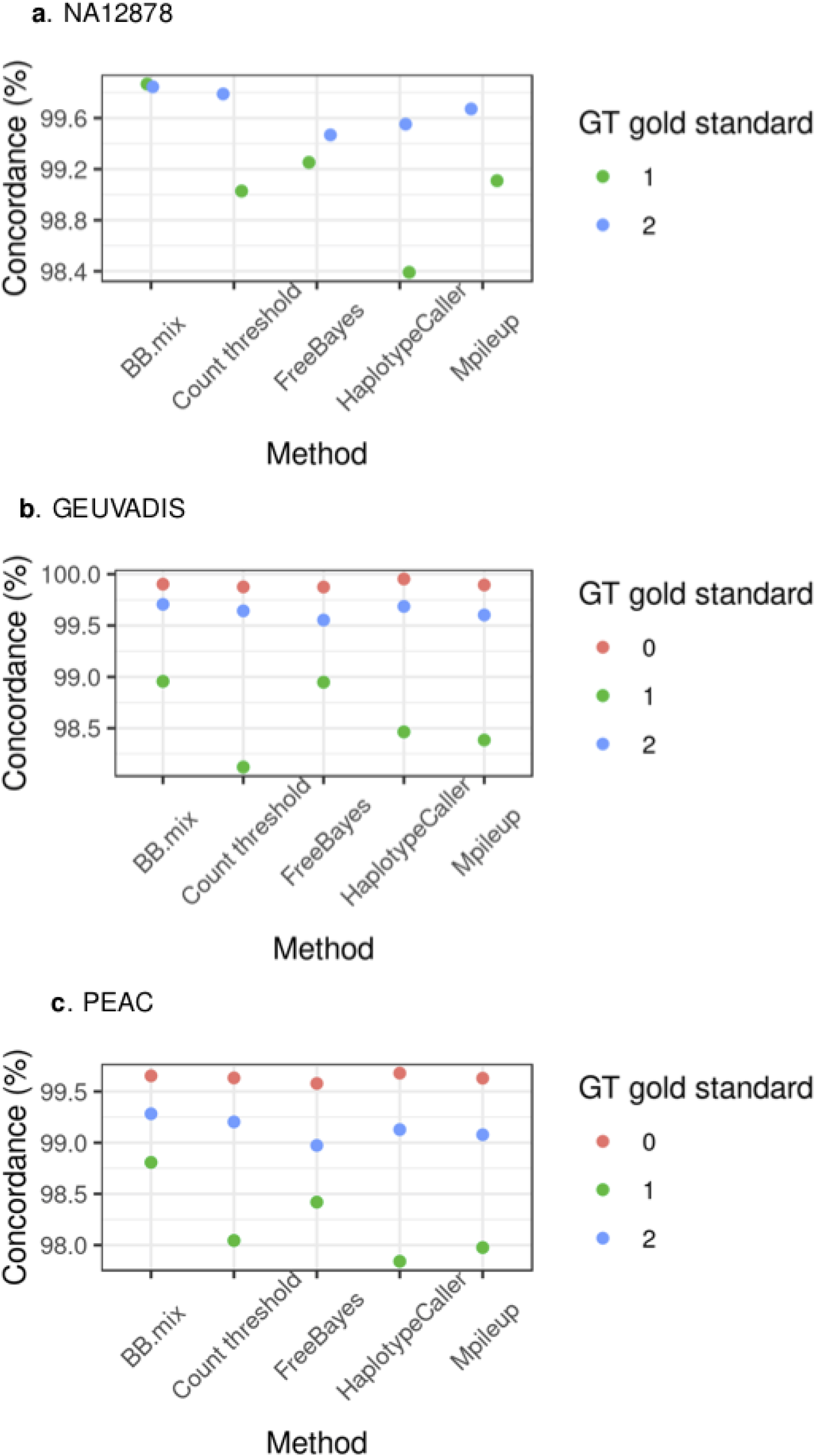
Concordance of calls by genotype. For each method the number of concordant calls relative to the gold standard for those SNPs with at least 10 supporting reads was calculated. The analysis was done for (a) Genome in a Bottle sample NA12878, (b) GEUVADIS samples and (c) PEAC samples. Note that for sample NA12878 the gold standard has only calls for heterozygous (1) and homozygous alternative (2) calls.

Next, we assessed the prevalence of homozygous calls in the gold standard called heterozygous by methods (hom to het errors) and vice versa (het to hom errors). Overall we observed a better performance of BBmix in both types of errors with the exception of GEUVADIS on hom to het errors with HaplotypeCaller performing better (**Fig. 3)**. FreeBayes was consistently worst for hom to het errors while HaplotypeCaller had the worst het to hom error rate for both NA12878 and PEAC (**Fig. 3**). Last, we compared methods performance across a wide range of read depth threshold on aggregated genotype calls. The accuracy of BBmix genotype calls was generally superior to competitor methods though the effect appeared modest especially for GEUVADIS and PEAC (**Supp. Fig. 2-4**). This is because roughly 60% of the calls were homozygous reference (Supp. Tables 1-3) which showed the most accuracy and least variability across methods, while for NA12878 the gold standard does not contain homozygous reference calls. As expected, increasing the depth threshold decreased the gap in accuracy between BBmix and Count threshold.

**Figure 3.**
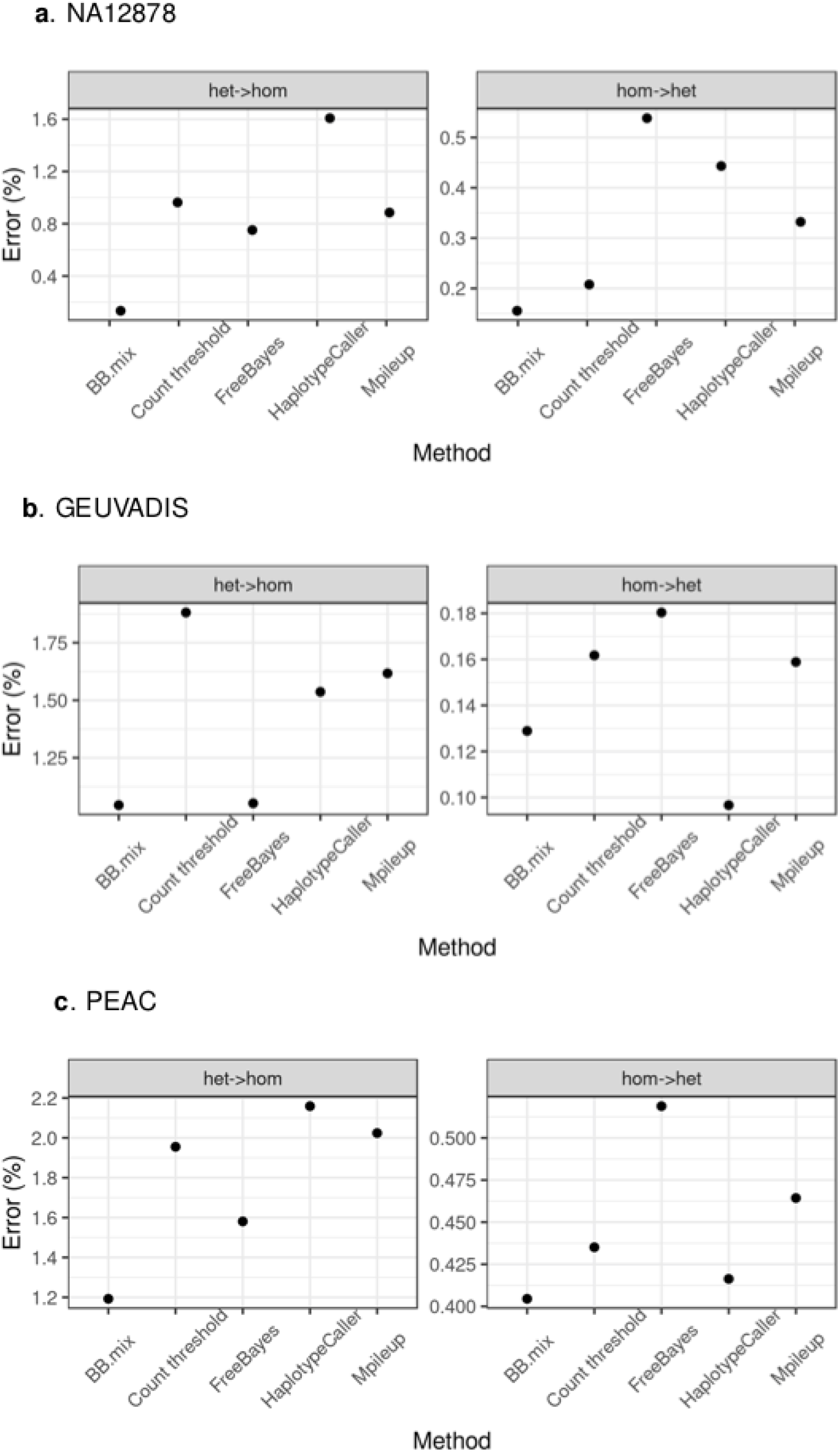
Genotype error type by method. For each method it is shown the percentage or erroneous calls for the indicated type (heterozygous in gold standard but homozygous by method (het -> hom) or vice versa). Only SNPs with at least 10 supporting reads were considered. (a) Genome in a Bottle sample NA12878, (b) GEUVADIS samples and (c) PEAC samples. Note that for sample NA12878 the gold standard has only calls for heterozygous (1) and homozygous alternative (2) calls.

With regards to the number of calls made by each method HaplotypeCaller had the highest number of calls, about 20-30% higher than BBmix, with the main gain being on homozygous reference calls (**Supp. Tables 1-3, Supp. Figs. 2b-4b**). For the subset of calls that were validated in the gold standard Mpileup had the highest number, between 10-25% higher than BBmix (**Supp. Figs. 2b-4b**). Overall BBmix accuracy was generally superior at the cost of lower number of calls.

### Exchangeability of model parameters between samples

Thus far we trained the BBmix model repeatedly for each sample under consideration. To save computational time we could train the model with a sample of reads from the pooled samples and use those parameters to call genotypes in each sample. Even better would be to have default parameters from an external sample and completely avoid model training. We assessed those options as follows: for sample NA12878, we trained the model with each of the 13 samples that were RNA-sequenced together with NA12878, with a pool of reads from all of the samples or using a randomly selected GEUVADIS external sample. We then called genotypes in NA12878 using the aforementioned scheme for model training and assessed genotype accuracy against the gold standard. The accuracy on genotypes was very robust to the source of reads for parameter training, ranging from 99.25%-99.60% for heterozygous calls and 99.40%-99.60% for homozygous alternative calls, and very close to the model trained with NA12878 reads, 99.38% for heterozygous and 99.44% for homozygous alternative calls (**Fig. 4a**). Next, we extended the analysis to the GEUVADIS sub-cohort by training the model with a pool of reads. We then called genotypes for each sample using the model trained either with the pool of reads, the external sample NA12878 or with the same sample used to fit the model. Again, the concordance of calls across all genotypes was very similar regardless of the source of reads used for training (**Fig. 4b**). Last, the same analysis performed in PEAC produced the same pattern, with very high concordance on genotype calls even when training with external samples (**Fig. 4c**).

**Figure 4.**
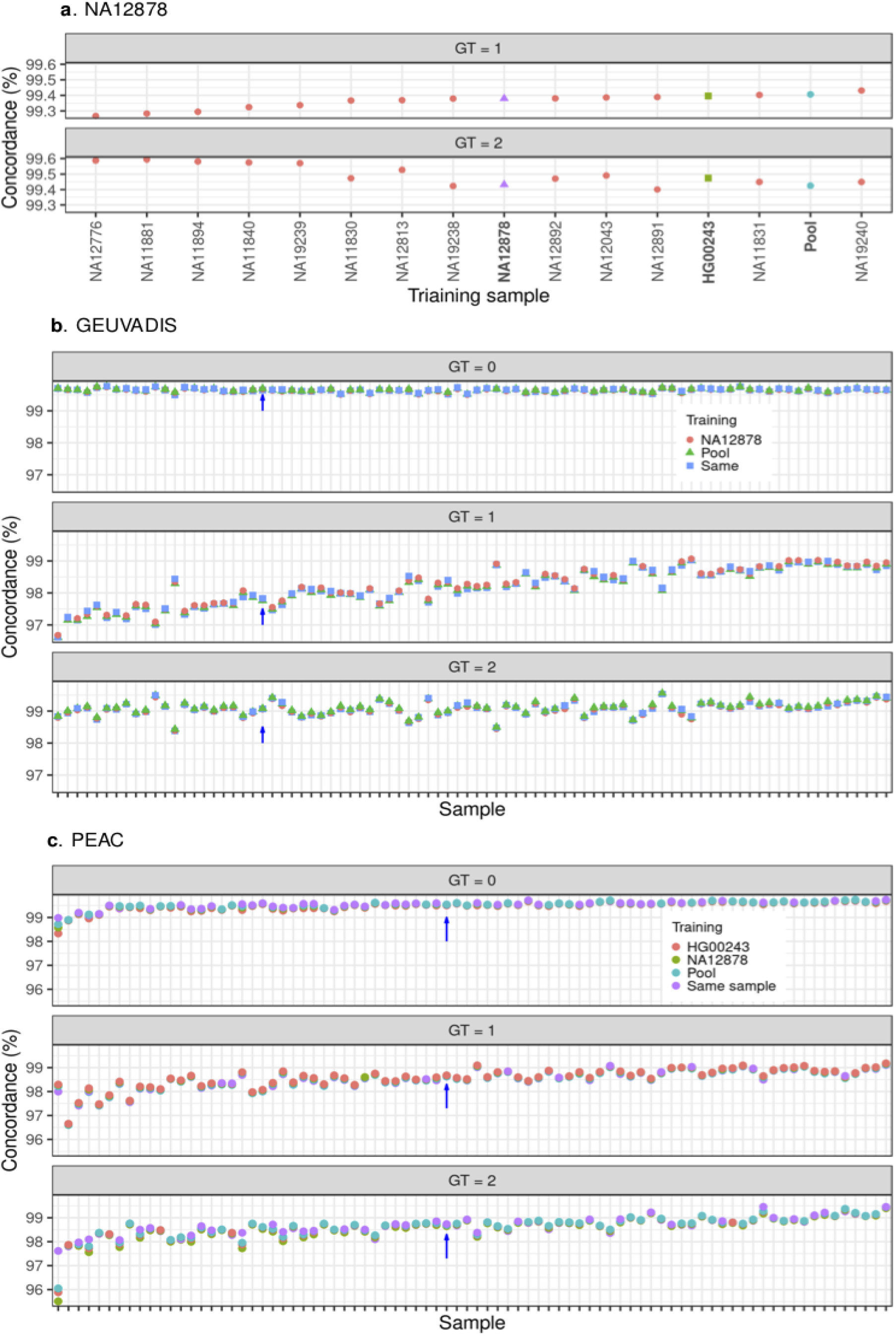
Exchangeability of model parameters. (a) The BBmix model was trained with each of the E-MTAB-1883 samples, a pool of those samples or a randomly selected external GEUVADIS sample (indicated in the x-axis). Genotypes were called in sample NA12878 and the concordance against the gold standard stratified by genotype is shown in the y-axis. (b) The BBmix model was trained with each of the GEUVADIS samples, a pool of them or the external sample NA12878. For each sample (x-axis) the plot shows the concordance on genotypes (y-axis) called with the same sample used for training the pool or NA12878. The arrow indicates sample HG00243 to ease comparison with **Supp. Fig. 6**. (c) Same as in (b) except that genotypes for the PEAC samples were additionally called using HG00243 as an external sample. The arrows indicate sample QMUL2009047 to ease comparison with **Supp. Fig. 6**.

Prompted by the high level of genotype concordance regardless of training sample, we compared the distribution of the posterior samples for the model parameters trained with either a pool of the respective study samples with NA12878, a representative sample for GEUVADIS or PEAC. In all examples we observed a mixed pattern with some parameters showing similar distributions but clear departures from the identity line as well (**Supp. Fig. 6**)

Overall, genotype calls appeared to be robust with regards to the source of reads used for model training.

## Discussion

Current pipelines for calling variants from RNA-seq rely on data cleaning and algorithms designed for DNA-seq, ignoring allele specific expression and reference mapping bias. In this study we provide proof of concept that calling variants from RNA-seq can be improved by modelling intrinsic sources of variation in RNA-seq read counts by assessing model performance in relation to DNA based methods for known common exonic SNPs and using DNA genotyping as the gold standard.

Our motivation to call variants from RNA-seq was to detect expression quantitative trait loci (eQTLs) using allele specific expression (ASE), which requires genotyping of exonic SNPs (Castel *et al*., 2015). This is because calling variants from RNA-seq has the potential to increase coverage for variants overlapping expressed genes. Moreover, genotyping errors arising from DNA-based genotyping may introduce bias, for example if homozygous exonic SNPs are mis-typed as heterozygous as the RNA-seq reads will show a strong allelic imbalance. We showed here that our method generally reduced this type of error compared to the other methods and performed particularly well on heterozygous calls, which is likely due to our modelling strategy accounting for noise in RNA-seq reads.

Our contribution shows that accounting for bias in RNA-seq can improve genotyping accuracy and we provide a statistical framework for modelling such data: in the case of the most carefully curated gold standard, the Genome in a Bottle sample, we reduced the proportion of erroneous calls over four-fold compared to the next best method. Although BBmix tends to increase accuracy, this occurs at the cost of reduced genotype calls made. Thus the method is designed to prioritise accuracy over volume. Although our method was not designed as a standalone pipeline for discovery, our modelling approach has potential to be integrated within current algorithms such as HaplotypeCaller, Mpileup or FreeBayes to tailor those methods for RNA-seq. Although Bayesian approaches tend to be computationally expensive, we showed that a modest number of training SNPs (1000) tends to be sufficient to learn the model parameters in less than an hour, and that calling variants was very consistent even when using an external sample, which completely eliminates the need for model training. Thus, our work presents a method tailored for calling variants using RNA-seq and also highlights that the current methods for calling variants using DNA could be adapted for RNA-seq by modelling RNA-seq count biases.

## Acknowledgements

This work was co-funded by the Wellcome Trust (WT220788), the MRC (MC_UU_ 00002/4) and supported by the NIHR Cambridge Biomedical Research Centre (BRC-1215-20014). M.L., A.B and C.P were supported by the MRC/Arthritis Research UK award: Maximising Therapeutic Utility in RA (MATURA) (Grant MR-K015346). The Pathobiology of Early Arthritis Cohort (PEAC) was supported by funding from the MRC (grant number G0800648).

