## Supplementary material for "BBmix: a Bayesian Beta-Binomial mixture model for accurate genotyping from RNA-sequencing"

### Supplementary Note 1: Beta-Binomial mixture statistical model

Calling variants with RNA-seq reads may lead to errors due to reference panel bias and allele specific expression, which may alter the ratio of allelic counts potentially favoring a homozygous call when a heterozygous is correct. To account for these sources of bias we developed a 2-step approach. First, we selected RNA-seq reads that overlap known exonic bi-allelic SNPs to learn the parameters for a mixture of three beta-binomial distributions underlying the three possible genotypes. In the second step we obtained posterior probabilities for each of the genotypes using the parameters learnt in step 1.

#### Fitting a Beta-Binomial mixture model to RNA-seq data

We selected RNA-seq reads overlapping a pre-defined set of feature SNPs (fSNPs) that we aim to genotype. We treat genotype as a latent variable, $G$, which corresponds to the number of alternative alleles at a given fSNP, $g = (0, 1, 2)$, and we assume that the distribution of reads differ according to its value. Given a fSNP $s$ indexed by $l$, we denote $R_{al}$ as the number of reads overlapping its alternative allele and $R_{tl}$ the total number of reads mapping $s_l$. We then fit a mixture of three Beta-Binomial distributions to the allelic counts of a random sample of $L$ fSNPs to estimate the parameters ($\mu$, vector with the mean of each component, and $\lambda$, vector with the overdispersion of each component) for the underlying distribution of genotypes. With $\theta$ a vector with the proportion of each component, we express the likelihood as:

$$\mathcal{L} = \prod_{l=1}^{l=L} \sum_{g=0}^{g=2} \theta_g BB(R_{al}|R_{tl}, \mu_g, \lambda_g)$$

where BB() is the beta binomial distribution:

$$f(R_{at}|R_{tl}, \alpha, \beta) = \frac{\binom{R_{al}}{R_{tl}} B(R_{al} + \alpha, R_{tl} - R_{al} + \beta)}{B(\alpha, \beta)}$$

and we reparameterize to: $\mu_g = \dfrac{\alpha}{\alpha + \beta}$ and $\lambda_g = \dfrac{1}{1 + \alpha + \beta}$

We put the following priors:

$\theta \sim Dirichlet(1, 1, 1)$
$\alpha = (1, 10, 499)$

$\beta = (499, 10, 1)$
$\mu_g \sim Beta(\alpha_g, \beta_g)$
$\lambda_g \sim \Gamma(\alpha_g + \beta_g, 1)$

We selected the Dirichlet prior as the natural choice for a set of probabilities that sum to 1, and the informative priors for $\alpha$ and $\beta$ were selected by visual inspection of the fit of beta distributions to the empirical distribution of reads for a collection of fSNPs with effector allele frequency (EAF) higher than 0.01 (Supp. Fig. 1).

We estimated $\boldsymbol{\mu}$ and $\boldsymbol{\lambda}$ by Markov chain Monte Carlo implemented in Stan (Stan development team, 2018). We checked convergence using Stan built-in n_eff and Rhat statistics as well as visual examination. We obtained 4000 posterior observations sampled from 4 chains for each parameter. For each chain we discarded the first 1000 observations.

### Calling genotypes

We call genotypes on a per-individual basis. For $s_l$ we calculated genotype likelihoods using posterior sample $j$ for each component of $\hat{\mu}$ and $\hat{\lambda}$ as follows:

$$P(R_{al}|R_{tl}, \hat{\mu}_{gj}, \hat{\lambda}_{gj}, G_l = g) = BB(R_{al}|R_{tl}, \hat{\mu}_{gj}, \hat{\lambda}_{gj}, G_l = g)$$

We derived priors for $s_l$ genotypes based on the estimated EAF ($\hat{f}$) obtained from the 1000G project phase 3 as:

$$P(G_l = 0) = (1 - \hat{f}_l)^2$$
$$P(G_l = 1) = 2 * (1 - \hat{f}_l) * (\hat{f}_l)$$
$$P(G_l = 2) = \hat{f}_l^2$$

To obtain posterior genotypes we use the $J = 4000$ posterior samples of $\mu_g$ and $\lambda_g$ as follows:

$$P(G_l = g|R_{al}, R_{tl}, \mu_j, \lambda_j) = \frac{1}{J} \sum_j \frac{P(G_l = g) * P(R_a|R_t, \mu_{gj}, \lambda_{gj}, G_l = g)}{\sum_{g=0,1,2} P(G_l = g) * P(R_a|R_t, \mu_{gj}, \lambda_{gj}, G_l = g)}$$

We defined the expected allele dosage for $s_l$ as $D_l$, so :

$$D_l = \frac{1}{J} \sum_j E[G_l|R_{al}, R_{tl}, \mu_j, \lambda_j] = \frac{1}{J} \sum_j D_{lj}$$

and $D_{lj}$ corresponds to:

$$D_{lj} = \begin{bmatrix} P(G_l = 1 | R_{al}, R_{tl}, \mu_{g=1,j}, \lambda_{g=1,j}) & P(G_l = 2 | R_{al}, R_{tl}, \mu_{g=2,j}, \lambda_{g=2,j}) \end{bmatrix} * \begin{bmatrix} 1 \\ 2 \end{bmatrix}$$

#### Genotype hard calling

We assign the most likely genotype if its probability is above a pre-defined threshold $p$, otherwise the genotype is set to missing.

### Supplementary Note 2: Data

The "Genome in a bottle" consortium has developed a pipeline integrating sequencing data generated by multiple technologies to produce a list of high confident heterozygous and homozygous alternative variant calls widely used for benchmarking and validation of variant calling pipelines https://www.nist.gov/programs-projects/genome-bottle. The sample employed is NA12878/HG001 from the HapMap project https://www.genome.gov/10001688/international-hapmap-project. Genotype calls were downloaded from "ftp://ftp-trace.ncbi.nlm.nih.gov/giab/ftp/release//NA12878_HG001/latest/GRCh37/HG001_GRCh37_GIAB_highconf_CG-IllFB-IllGATKHC-Ion-10X-SOLID_CHROM1-X_v.3.3.2_highconf_PGandRTGphasetransfer.vcf.gz". In addition, RNA-seq data produced from a lymphoblastic cell line derived from sample NA12878 in addition to 13 other samples from the 1000 Genome project were available in the study E-MTAB-1883 at array express https://www.ebi.ac.uk/arrayexpress/experiments/E-MTAB-1883/.

We downloaded RNA-seq data from 86 GEUVADIS lymphoblastic cell line samples with EUR ancestry (GBR code) from ArrayExpress (E-GEUV-1, Supplementary Table 4). Genotypes by DNA-sequencing are publicly available from the 1000 Genome consortium https://www.internationalgenome.org/data-portal/data-collection at https://www.internationalgenome.org/data-portal/data-collection/phase-3.

For our "real data" example we used RNA-seq samples from synovial tissue for 82 samples of the Pathobiology of Early Arthritis Cohort (PEAC) study(https://www.ncbi.nlm.nih.gov/pmc/articles/PMC6718830/#mmc1 available at array express (E-MTAB-6141) https://www.ebi.ac.uk/arrayexpress/experiments/E-MTAB-6141/samples/DNA-microarray. Genotypes by DNA-microarray are available upon reasonable request.

The file used to exclude difficult to map regions in the chromosome was downloaded from phASER https://github.com/secastel/phaser/tree/master/phaser at https://www.dropbox.com/s/fbfntaa4oc75x6m/hg19_hla.bed.gz?dl=0

### Supplementary Note 3: RNA-seq processing

Starting from fastq files the following steps were performed following the GATK best practices pipeline. Details on the implementation of each step can be found at `https://gitlab.com/evigorito/bbmix_pipeline/-/blob/master/Snakefile`

#### Alignment

All samples were aligned to the human genome assembly GRCh37 using 2-pass STAR (DOI: 10.1002/0471250953.bi1114s51 ).

#### Removal of duplicate reads

Removal of duplicate reads which may arise from PCR artifacts was performed using the implementation by WASP (10.1038/nmeth.3582) coded in rmdup_pe.py `https://github.com/bmvdgeijn/WASP`. This implementation removes reads at random with regards to mapping score in order to minimise the risk of reference mapping bias.

#### Base quality re-calibration

This steps applies machine learning to detect and correct for patterns of systematic errors in the base quality scores of the alignments.

### Supplementary Note 4: Comparing genotyping callers

Details for the implementation of HaplotypeCaller, Mpileup and FreeBayes can be found in the Snakefile at `https://gitlab.com/evigorito/bbmixpaper_pipeline`. For the BBmix method refer to the Snakefile at `https://gitlab.com/evigorito/bbmix_pipeline`

#### HaplotypeCaller

HaplotypeCaller is the algorithm designed for calling genotypes in the GATK pieline, which we applied with default parameters. The resulting calls were filtered using the recommended annotations internally generated by HaplotypeCaller with default parameters. Briefly, SNPs were excluded if they clustered with at least 3 SNPs within a 35 bp window, if they showed evidence of strand bias (SNPs with a p-value of a Fisher exact test Phred scaled over 30) or their quality adjusted by depth was lower than 2. The selected annotations and values are the default recommended at (`http://barcwiki.wi.mit.edu/wiki/SOP/CallingVariantsRNAseq`)

#### Mpileup/bcftools

For calling SNPs we fed the aligned files obtained after base quality re-calibration step on the GATK pipeline into bcftools version 8 using the "mpileup" and "call" commands selecting reads with a quality score of at least 20 and using default parameters.

#### FreeBayes

The recommended input for FreeBayes are aligned files with PCR duplicates removed (`http://clavius.bc.edu/~erik/CSHL-advanced-sequencing/freebayes-tutorial.html`. FreeBayes was run with default parameters.

#### BBmix model

We used the aligned files obtained after base quality re-calibration following the GATK pipeline to count uniquely mapped reads overlapping the set of SNPs we wanted to genotype (section Selecting exonic SNPs), implemented in a Python script. The script also detects SNPs in clusters using the same parameters as in the GATK pipeline (3 SNPs within 35 bp). We do not model base quality, instead we impose a threshold of 20 and we select uniquely mapped reads. After parameter training and calling variants (statistical model section), we discarded variants with evidence of strand bias or in clusters following the same thresholds recommended in the GATK pipeline. We implemented those filters in Python. We also excluded SNPs with no evidence of variation (homozygous in all samples), similar to other methods.

### Count threshold

We implemented this method as a direct comparison to the BBmix model. In this case we called hard genotypes based on the proportion of reads mapping the alternative allele ($p_l = \frac{R_{al}}{R_{tl}}$) as follows:

$p_l \leq 0.1$: Homozygous reference

$0.1 < p_l < 0.9$: Heterozygous

$p_l \geq 0.9$: Homozygous alternative

We refer to this method as "Count threshold". This way we were able to directly assess the benefit of implementing the statistical model relative to calling genotypes just based on intuitive thresholds. We applied the same filters as when using the BBmix model.

### Selecting exonic SNPs

Exonic SNPs with a minor allele frequency higher than 0.01 were selected as those overlapping the union of exons which are annotated in the 1000 Genome phase3 legend files. HLA and regions known for allele mapping bias were excluded from the analysis (DOI:https://doi.org/10.1038/ncomms12817, phASER files from `https://github.com/secastel/phaser/tree/master/phaser`).

### Assessing model performance

For each dataset we considered the DNA-based genotyping as the gold standard. Concordance was calculated as:

$$\text{Concordance} = 1 - \sum_l (D_l - G_l)^2 / L$$

$D_l$ dosage for$SNP_l$ obtained by method

$G_l$ genotype for $SNP_l$ in the gold standard

Alternatively, for hard calls we calculated the proportion of calls that were correctly genotyped by a method relative to the gold standard.

### Supplementary Figures

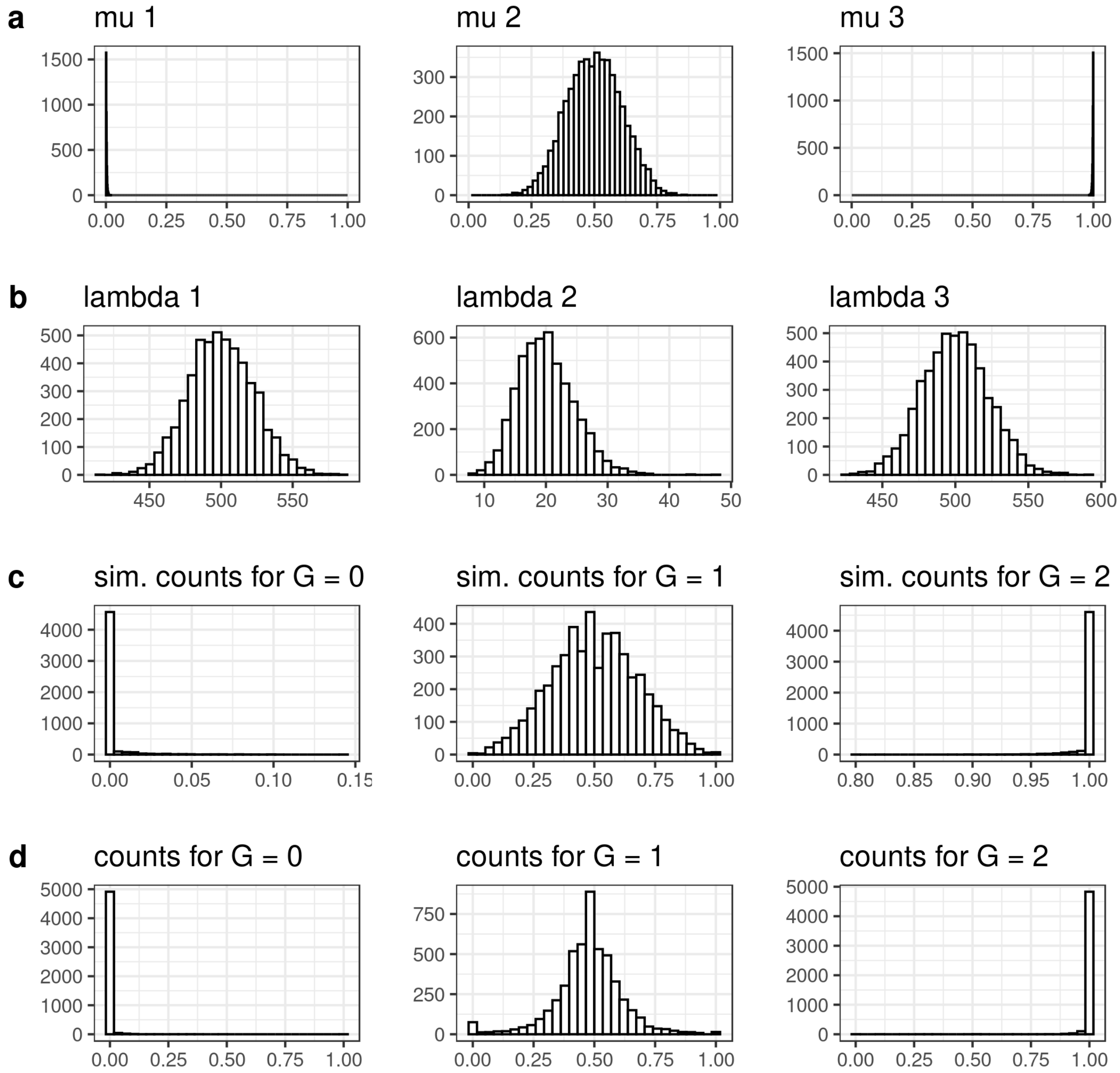


Supplementary Figure 1: Priors for BBmix model.

a) Distribution of the three mean components of the model, mu1-mu3 generated from 5,000 draws from a Beta distribution with the $\alpha$ and $\beta$ parameters [1,10,499] and [499,10,1], respectively. (b) Distribution of the three components of the over-dispersion parameter (lambda1-3) generated from 5,000 draws from a $\Gamma(\alpha+\beta, 1)$ distribution. (c) Distribution of the simulated proportion of reads mapping alternative alleles conditional on the total number of reads. The proportion of reads overlapping the alternative allele for each component (p1-p3) was simulated from 5000 draws from a Beta distribution with mean $\mu$ generated in (a) and over-dispersion $\lambda$ generated in (b). Then, the reads overlapping the alternative allele were simulated from 5000 draws of a Binomial distribution with total reads from a random sample of observed reads from 5000 SNPs of a GEUVADIS sample with at least 10 reads supporting each SNP and probability p1-p3. Last, the proportion of reads overlapping the alternative allele was calculated as the ratio between the simulated number of reads overlapping the alternative allele and the observed total number of reads. (d) For comparison the empirical proportion of reads overlapping the alternative allele obtained using raw counts from a GEUVADIS sample (HG002215) and stratified by the genotype in the gold standard. For each genotype the proportion of reads corresponding to 5000 SNPs are shown. labelfig:priors

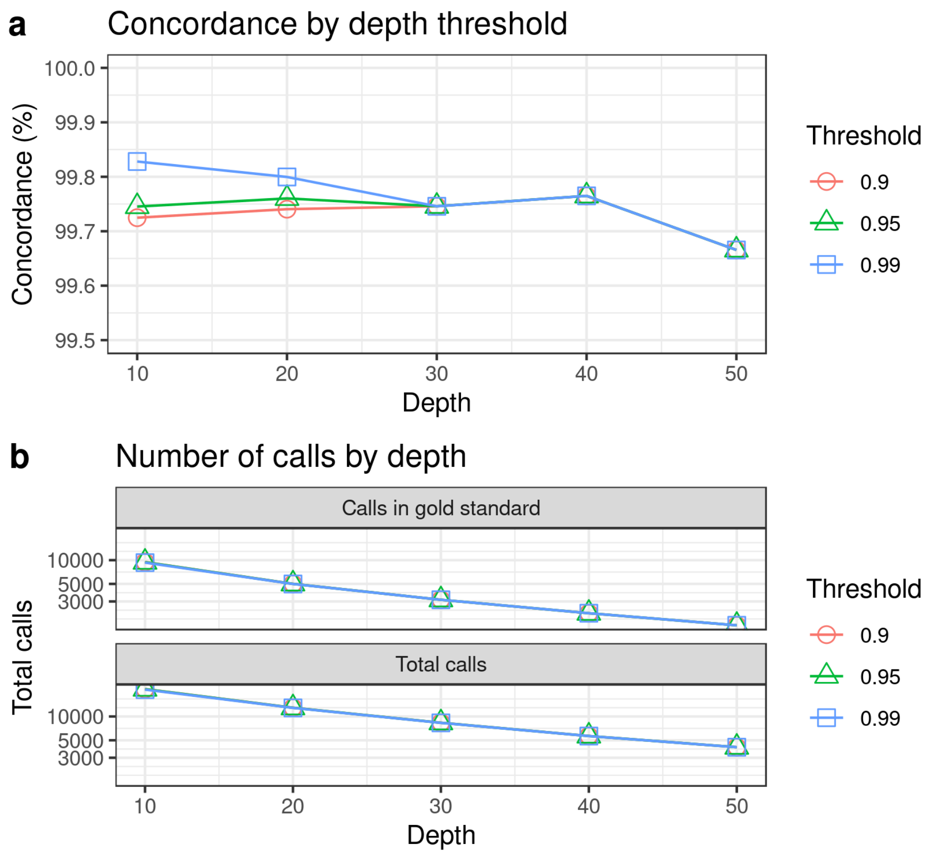


Supplementary Figure 2: Hard thresholds for Beta-Binomial mixture model. We made hard calls for sample NA12878 using thresholds 0.9, 0.95 and 0.99. (a) Concordance against the gold standard using high confident calls at the indicated depth. (b) Number of calls at the indicated depth for hard calls at the indicated thresholds (0.9, 0.95 and 0.99).

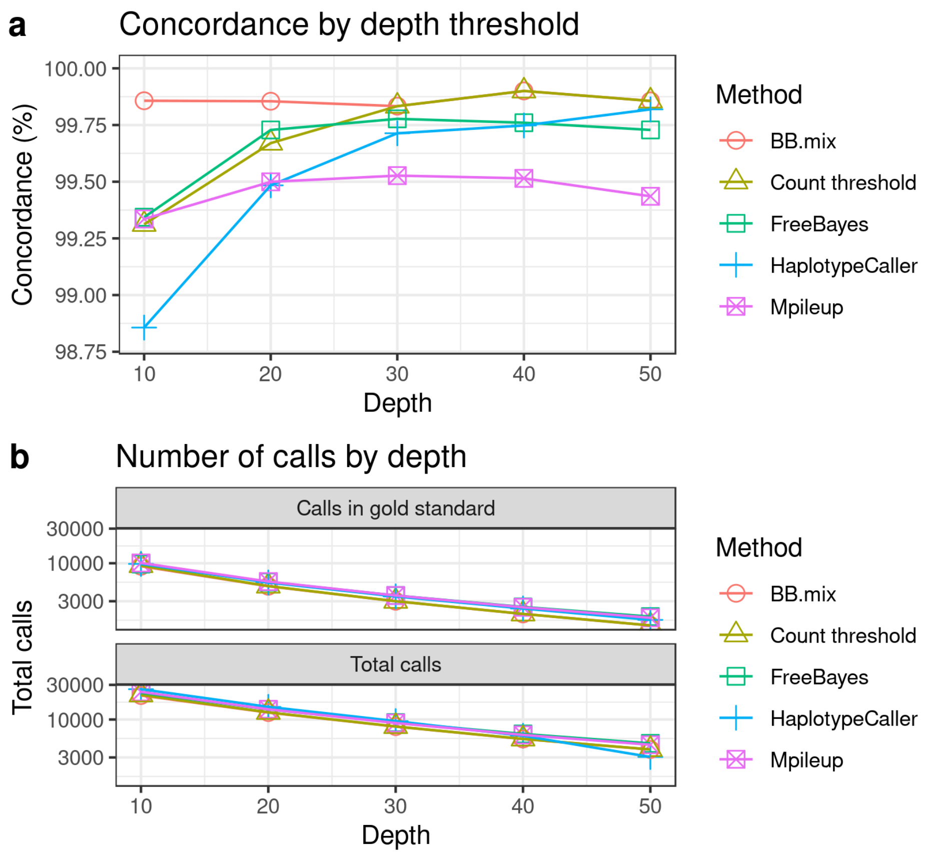


Supplementary Figure 3: Benchmarking methods using Genome in a Bottle sample NA12878. Genotypes were called using RNA-seq with FreeBayes, HaplotypeCaller, Mpileup, BBmix and Count threshold. For each method we selected calls with at least 10, 20, 30, 40 or 50 supporting reads, as defined by each method. (a) Concordance was evaluated on hard calls against the set of high confidence genotype calls publicly available for this sample. BBmix and Count threshold lines overlapped for depth thresholds of 40 and 50. (b) The solid lines correspond to the total number of calls made by each method by depth. The dashed lines correspond to the total number of calls that could be validated against the gold standard (ie sites that had a genotype call in the gold standard). For this sample the gold standard has only homozygous alternative and heterozygous calls.

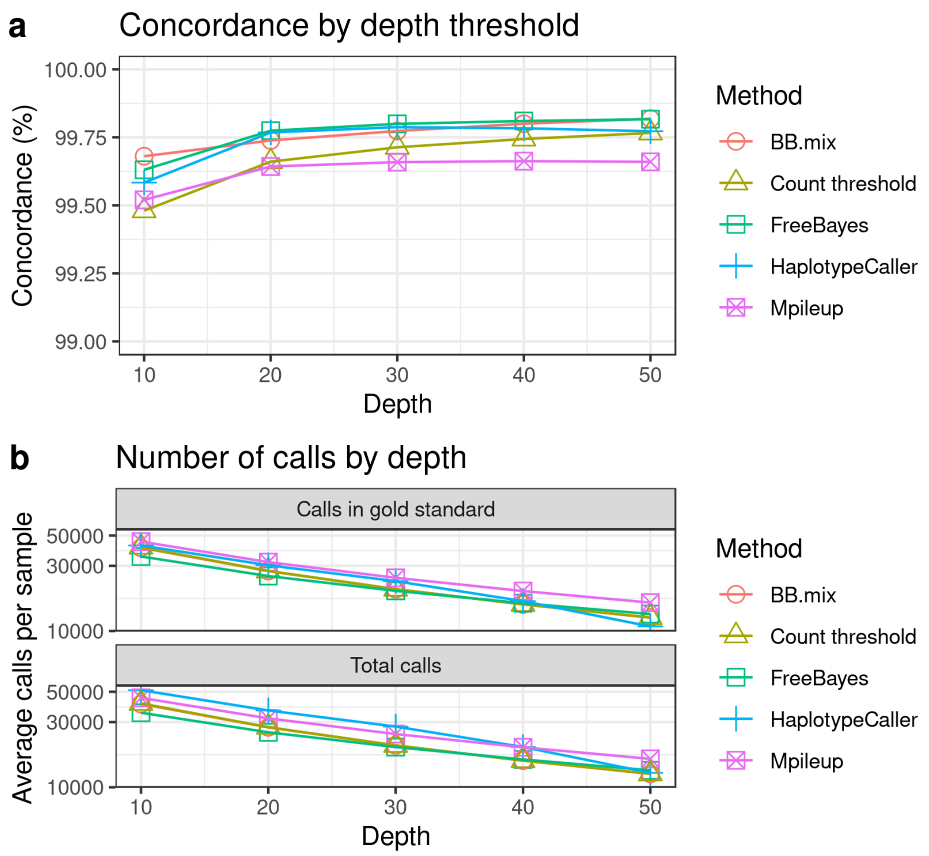


Supplementary Figure 4: Benchmarking methods using GBR GEUVADIS samples. Genotypes were called using RNA-seq with FreeBayes, HaplotypeCaller, Mpileup, our Beta-Binomial mixture model (BB.mix) and Count threshold. For each method we selected calls with at least 10, 20, 30, 40 or 50 supporting reads. Each method was run using default parameters. (a) Concordance was evaluated on hard calls against DNA-seq calls publicly available in the 1000G phase 3 project. (b) Average number of calls per sample for those validated with the gold standard (top) or total calls (bottom).

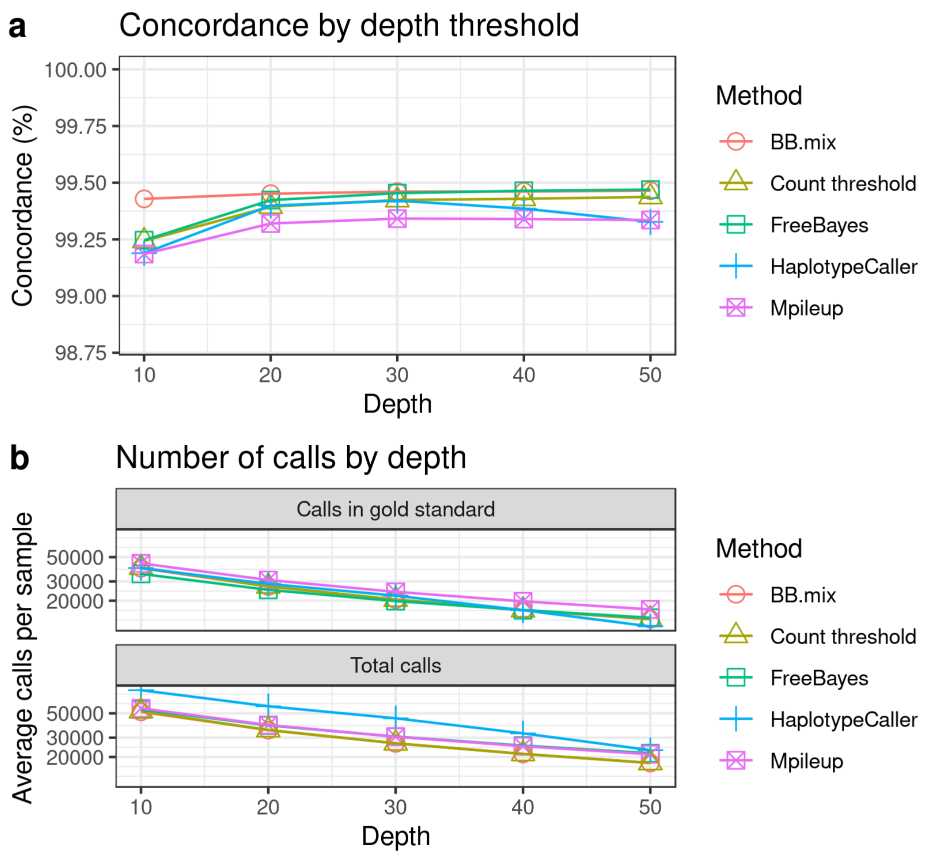


Supplementary Figure 5: Benchmarking methods using PEAC samples. Genotypes were called using RNA-seq with FreeBayes, HaplotypeCaller, Mpileup, BBmix and Count threshold. For the gold standard (genotyping with DNA-microarrays) we only considered calls with probability 1. For each method we selected calls with at least 10, 20, 30, 40 or 50 supporting reads. Each method was run using default parameters. (a) Concordance was evaluated on hard calls against DNA-microarray calls (Methods section). (b) Average number of calls per sample for those validated with the gold standard (top) or total calls (bottom).

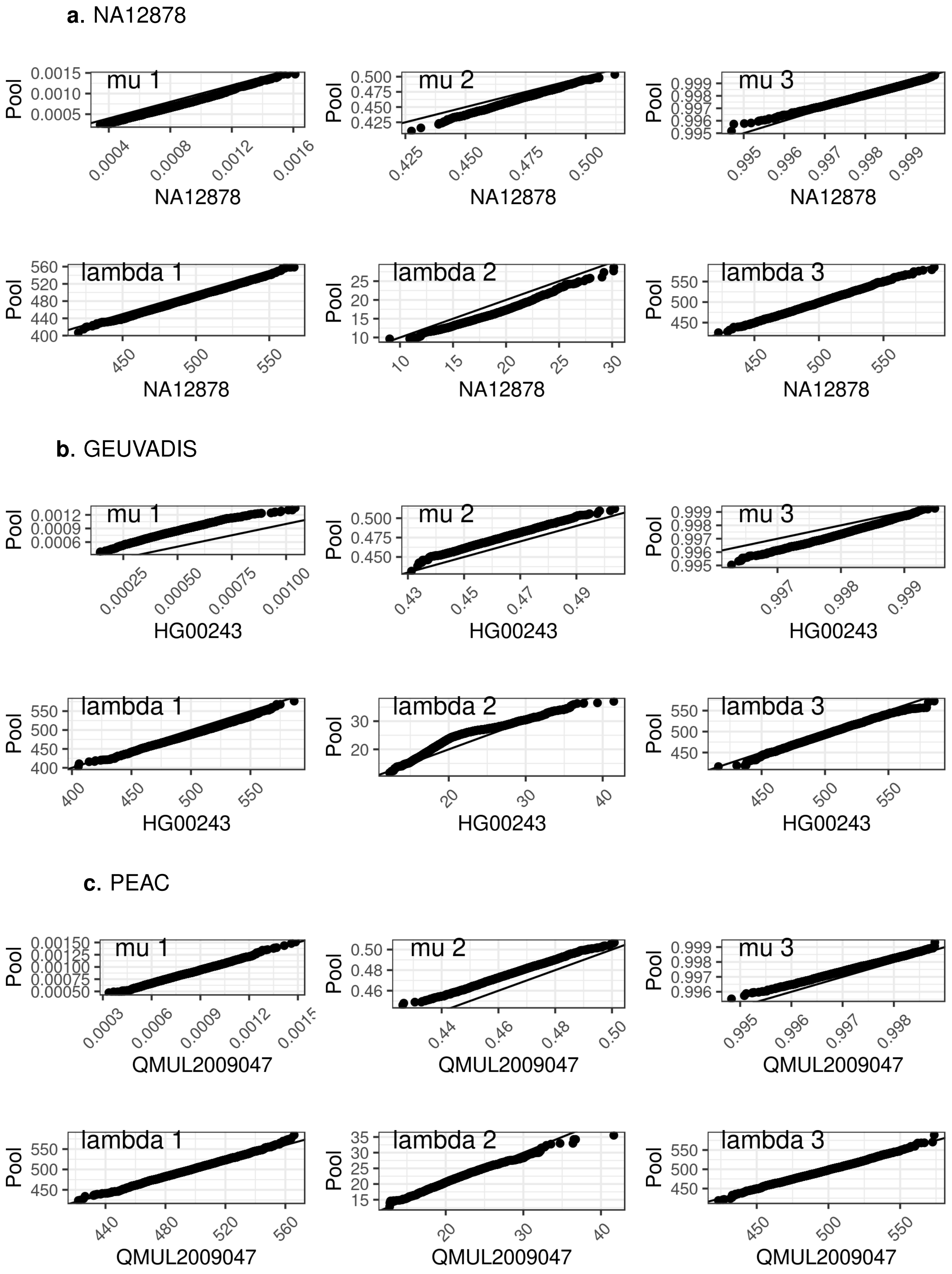


Supplementary Figure 6: Exchangeability of model parameters between samples.

The QQ plot shows posterior samples (4000) for each of the model parameters trained with the pool of study samples or the indicated sample. The y=x line is shown in black. (a) NA12878, (b) GEUVADIS, (c) PEAC.

### Supplementary Tables

| Method | Genotype 0 | 1 | 2 | Total |
|---|---|---|---|---|
| BBmix | 11641 | 3504 | 6265 | 21410 |
| Count threshold | 11891 | 3665 | 6313 | 21869 |
| FreeBayes | 12843 | 4017 | 6426 | 23286 |
| HaplotypeCaller | 15569 | 4092 | 6498 | 26159 |
| Mpileup | 13231 | 4189 | 6590 | 24010 |

Supplementary Table 1: Number of calls made by each method stratified by genotype for sample NA12878.

| Method | Genotype 0 | 1 | 2 | Total |
|---|---|---|---|---|
| BBmix | 27126 ± 4039 | 7913 ± 1468 | 5213 ± 910 | 40252 ± 6246 |
| Count threshold | 27434 ± 4008 | 8472 ± 1301 | 5309 ± 916 | 41215 ± 6204 |
| FreeBayes | 22818 ± 2754 | 7576 ± 1013 | 4724 ± 675 | 35119 ± 4413 |
| HaplotypeCaller | 35634 ± 4703 | 9555 ± 1279 | 6172 ± 869 | 51361 ± 6823 |
| Mpileup | 28852 ± 3645 | 9821 ± 1332 | 6460 ± 940 | 45134 ± 5889 |

Supplementary Table 2: Mean and standard deviation of the number of calls per sample made by each method stratified by genotype using GBR-GEUVADIS samples.

| Method | Genotype 0 | 1 | 2 | Total |
|---|---|---|---|---|
| BBmix | 34204 ± 11998 | 10491 ± 3877 | 6893 ± 2658 | 51588 ± 18448 |
| Count threshold | 34611 ± 12028 | 10567 ± 3887 | 7117 ± 2683 | 52295 ± 18510 |
| FreeBayes | 34767 ± 8586 | 11963 ± 3147 | 7521 ± 2075 | 54251 ± 13714 |
| HaplotypeCaller | 61969 ± 19368 | 11752 ± 3786 | 7605 ± 2486 | 81326 ± 25524 |
| Mpileup | 35156 ± 11003 | 12152 ± 4043 | 8237 ± 2787 | 55545 ± 17722 |

Supplementary Table 3: Mean and standard deviation of the number of calls per sample made by each method stratified by genotype using samples from the PEAC study.
